# Genotyping-by-sequencing supports a genetic basis for alpine wing-reduction in a New Zealand stonefly

**DOI:** 10.1101/264473

**Authors:** Andrew J. Veale, Brodie J. Foster, Peter K. Dearden, Jonathan M. Waters

**Affiliations:** Department of Zoology, University of Otago, Dunedin 9016, New Zealand; Department of Biochemistry, University of Otago, Dunedin 9016, New Zealand

## Abstract

Wing polymorphism is a prominent feature of numerous insect groups, but the genomic basis for this diversity remains poorly understood. Wing reduction is a commonly observed trait in many species of stoneflies, particularly in cold or alpine environments. The widespread New Zealand stonefly *Zelandoperla fenestrata* species group (*Z. fenestrata, Z. tillyardi, Z. pennulata*) contains populations ranging from long-winged (macropterous) to vestigial-winged (micropterous), with the latter phenotype typically associated with high altitudes. The presence of flightless forms on numerous mountain ranges, separated by lowland fully winged populations, suggests wing reduction has occurred multiple times. We use Genotyping by Sequencing (GBS) to test for genetic differentiation between fully winged (n=62) and vestigial-winged (n=34) individuals, sampled from a sympatric population of distinct wing morphotypes, to test for a genetic basis for wing morphology. We found no population genetic differentiation between these two morphotypes across 6,843 SNP loci, however we did detect several outlier loci that strongly differentiated morphotypes across independent tests. This indicates small regions of the genome are likely to be highly differentiated between morphotypes, indicating a genetic basis for morphotype differentiation. These results provide a clear basis for ongoing genomic analysis to elucidate critical regulatory pathways for wing development in Pterygota.

## Introduction

Understanding the genetic basis of phenotypic variability not only illuminates the active evolutionary processes occurring within species but may also shed light on the evolution of different morphologies among species. Wing polymorphism has arisen in many insect orders, with variability in wing morphology prominent in Hemiptera (true bugs), Coleoptera (beetles), Orthoptera (crickets and grasshoppers), and Plecoptera (stoneflies) ^1-4^. Within these groups, species that have lost flight are particularly common on islands, at high altitudes and high latitudes ^1^. The degree of wing development may vary between closely related species or within a species. While referred to as “wing polymorphism”, this variation often consists of morphs that differ in all major aspects of flight capability (e.g. size of flight muscles, production of flight fuels), as well as many other aspects of physiology and reproduction. These polymorphisms may result from a variety of causes: alternate morphologies may be encoded by different genotypes (genetic polymorphism), induced by different environments (environmental polyphenism), or produced by variation in both genetic and environmental factors ^5^. The degree of wing development can either be dimorphic with two alternative forms, or variation can exist along a spectrum.

There are many factors that influence the relative costs and benefits of flight in insects (reviewed by ^2,6-8^). Wing reduction may confer an adaptive advantage when habitat stability is high, and when habitat complexity is low ^9^. Habitat isolation may also promote flight loss, as the removal of flighted emigrants from habitat patches selects against this dispersal ability ^7,10-12 1^. Specifically, in alpine environments high winds may sweep away individuals with long wings ^7,13-15^. Wing reduction has also been attributed to the high energy expenditure required in the production and maintenance of flight apparatus, which are traded off at the expense of other life-history traits – particularly fecundity ^1,4,16-21^.

Stoneflies are of particular interest relating to the evolution of insect flight because of their early divergence within winged insects (Pterygota) and since they exhibit multiple wing-powered locomotive behaviors, including sailing and skimming on the water surface ^22^. These methods of locomotion have even been proposed as models for the evolution of flight in insects ^23-25^, and it has been suggested that stoneflies thus may exhibit an ancestral form of wing and flight development ^22,26^. Many stonefly species have reduced wings, with four forms of wing-length polymorphism described: macropterism (fully winged or long-winged), brachypterism (short-winged), micropterism (vestigial-winged) and apterism (wingless) ^27^. Even fully winged stonefly taxa are typically considered to be weak flyers with limited dispersal ability ^27-33^. There have been several studies of wing reduction in stoneflies e.g. ^13,15,32,34-38^, with some suggesting a possible genetic basis for short wingedness e.g. ^39^ but this hypothesis remains to be tested.

Over the last decade, high-throughput genetic sequencing, along with reduced representation genomic libraries ^40^ have enabled the low-cost discovery and genotyping of thousands of genetic markers for non-model organisms, revolutionizing ecological, evolutionary and conservation genetics ^41-43^. In particular, these advances have enabled the discovery of many candidate loci involved in specific phenotypic traits ^44-46^. Such advances have been made either with quantitative trait loci (QTL) mapping using pedigree information, or through genome-wide association studies (GWAS) that identify non-random associations of alleles between loci and adaptive traits as a consequence of natural selection ^47-49^.

The underlying bases for wing polymorphism have now been studied in several species of insects, showing various environmental, developmental, and genetic controls, often with multiple developmental pathways and regulators e.g. ^50^. For instance, the proximate endocrine processes that control wing development have been investigated in wing-polymorphic crickets *(Gryllus sp.),* showing Juvenile Hormone (JH) may regulate wing development in this species ^5,51^, while in a planthopper *(Nilaparvata lugensor),* genes in the insulin-signaling pathway may regulate wing development ^52,53^. The genes responsible for wing polymorphism have also recently been investigated in ants *(Pheidole morrisi)* ^54^, salt marsh beetles *(Pogonus chalceus)* ^55^, and pea aphids *(Acyrthosiphon pisum)* ^56,57^. There are also known genes responsible for wing patterning and development in model organisms such as *Drosophila melanogaster,* which may be relevant to intra-specific wing polymorphism ^58^. While genetic changes often underlie wing polymorphism,epigenetic changes have also been demonstrated between wing morphs in a planthopper *(Sogatella furcifera)* ^59,60^.

The New Zealand stonefly *Zelandoperla fenestrata* species group *(Z. fenestrata, Z. pennulata, Z. tillyardi)* contains populations that range from fully winged to vestigial-winged, with wing-reduced populations more prevalent in southern South Island, particularly at higher altitudes ^61,62^. Under current taxonomy micropterous individuals are classified as *Zelandoperla pennulata* (McLellan 1967), dark-colored individuals including those implicated in the mimicry of another stonefly *(Austroperla cyrene)* are classified as *Zelandoperla tillyardi* (McLellan 1999), while the remaining light-colored fully winged individuals are classified as *Zelandoperla fenestrata* (Tillyard 1923). The three described species, however, appear to represent co-distributed color and wing-length polymorphisms rather than discrete evolutionary units, with the species group actually comprising five geographically discrete, deeply divergent clades (from 2% - 9% average divergence at COI) ^32^. These five regional clades exhibit differing propensities to exhibit wing reduced populations. Of the five clades of *Z. fenestrata* species group, Clade 1 is generally wing-dimorphic, with fully winged lowland populations and alpine associated vestigial-winged populations, with a steep transition in wing morphology at around 500 m.a.s.l (Figure 1). In contrast, Clades 2-4 appear to be comprised of only fully winged individuals, and Clade 5 is thought to be exclusively micropterous or apterous ^62^. Given the level of divergence between clades, and the probable differences in developmental characteristics between them, these clades may represent different species; further study is warranted to reclassify this group. The believed difference in propensity for wing reduction in different clades may suggest the possibility of a genetic basis for wing reduction in these taxa. Furthermore, the presence of non-dispersive, flightless forms on multiple mountain ranges in *Z. fenestrata* Clade 1, separated by lowland winged populations, suggests wing reduction may have evolved multiple times in this lineage ^32^. At finer spatial scales, recent genetic studies have shown phylogenetic divergence in wing-reduced populations of *Z. fenestrata* Clade 1 between adjacent mountain streams, highlighting the low dispersal ability of alpine populations and the possibility that each stream may have been colonized independently by winged lowland ancestors ^63^. The specific mechanisms and genes behind wing development and polymorphism in *Z. fenestrata* remain unknown.

**Figure 1.**
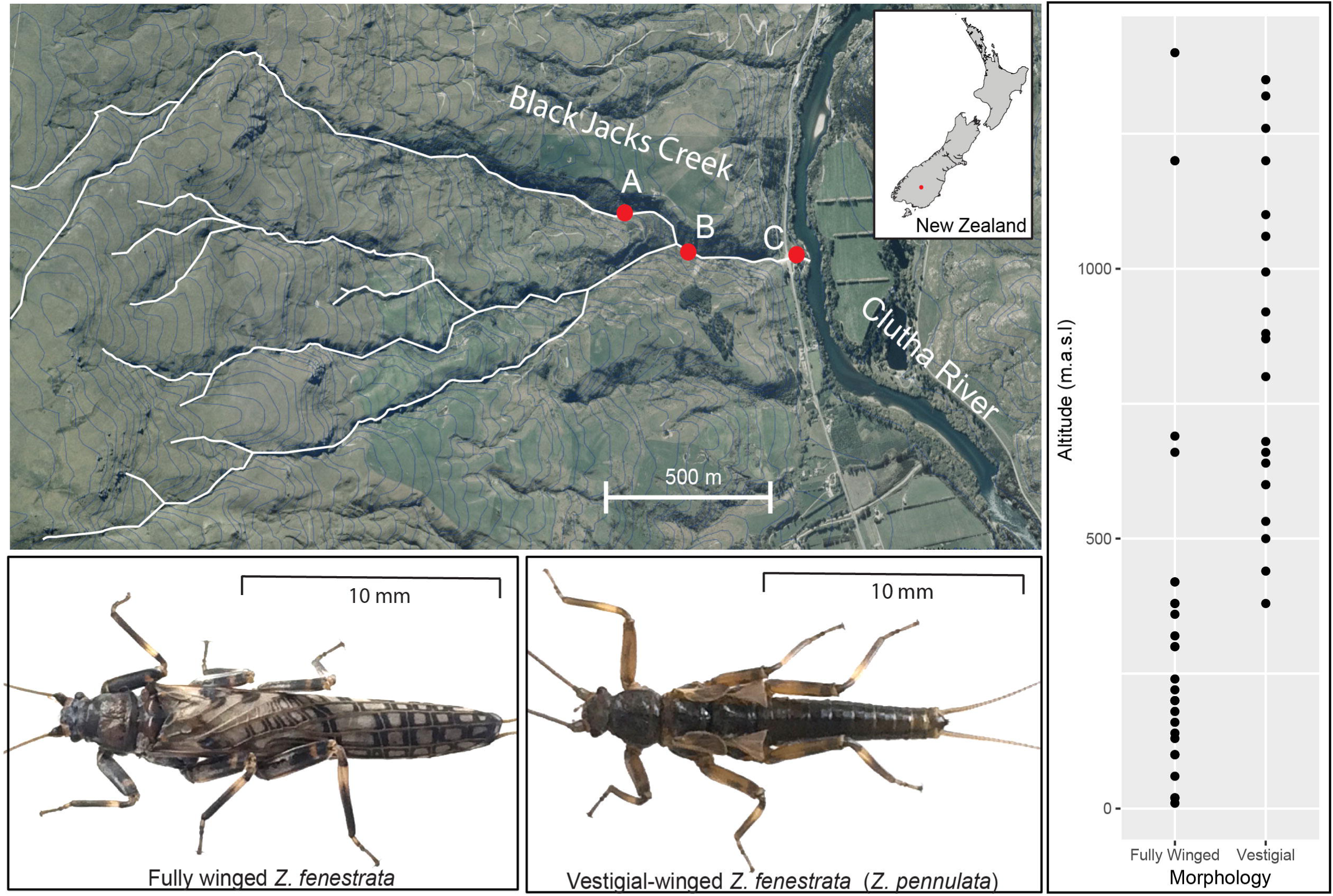
Map showing the sampling locations along Black Jacks Creek (A = 200 m.a.s.l, B = 130 m.a.s.l, C = 90 m.a.s.l). Inset below are examples of the two morphotypes to scale. To the right are the regional patterns of fully winged and vestigial-winged *Z. fenestrata* Clade 1 (data from McCulloch et al., 2009).

There are two (non-exclusive) hypotheses as to how *Z. fenestrata* Clade 1 lose their wings: 1) wing loss is genetically determined, or 2) wing loss is mediated by environmentally determined gene expression (i.e. polyphenism). Both of these hypotheses have received support from studies of other wing-dimorphic insects. Examples of taxa showing genetically determined wing dimorphism (Hypothesis 1) include several species of carabids and weevils ^14,64,65^ where wing dimorphism is controlled by a single gene operating in a Mendelian fashion. Similarly, in field crickets ^66^ and maize leaf hoppers *(Cicadulina sp.)* ^67^, wing polymorphism is genetically controlled but related to a complex interplay between many genes. However, in a situation more consistent with Hypothesis 2 (polyphenism), while wing morphology in *Gryllus* crickets can be controlled either by a single gene locus or a polygene complex, both can be regulated by the level of juvenile hormone (JH) – whereby if JH exceeds a threshold value during a critical developmental stage of the insect, wing development is suppressed ^5,51,68^. Other environmental factors that can influence wing development include abiotic factors such as temperature ^65^ and photoperiod ^69^ as well as biotic factors such as food resources ^65^ and population density ^70^. Many of these environmental regulators of wing development also have a genetic component, for instance the fully winged morphotype of the red fire bug *(Pyrrhocoris apterus)* is determined by a recessive allele, whose penetrance depends on photoperiod and temperature ^71^. Environmentally induced wing polyphenism in insects can also be transgenerational, with the level of the hormone ecdysone in the mother (regulated by population density) altering the expression of wing development in the offspring of the pea aphid *(Acyrthosiphon pisum)* ^72^.

In this study, we use Genotyping By Sequencing (GBS) to test for genetic differentiation between wing morphotypes in *Z. fenestrata* Clade 1, and test for loci specifically associated with wing reduction. GBS analyses a subset of the genome next to specific restriction sites, providing a near random sample of SNP loci across the genome, some of which may be associated with differentially adaptive genes or regulatory regions ^47-49^. As mentioned, *Z. fenestrata* Clade 1 is a divergent clade of the species group, with a propensity for alpine related wing-reduction, and it may be divergent enough to other clades to warrant reclassification to species or sub-species level. Surveys of *Z. fenestrata* Clade 1 morphotype distributions conducted by our lab identified one stream (Black Jacks Creek) that exhibited an unusual pattern of high overlap between wing morphotype populations at a low altitude. By focusing our study on a single stream population that exhibits co-distributed extreme wing morphologies, we aim to examine genomic differentiation between morphotypes without the confounding factor of neutral genetic population structure or other environmental differences.

## Methods

### SAMPLE COLLECTION

Sampling was conducted along Black Jacks Creek (on the Old Man Range, South Island, New Zealand, at three sampling zones (80 – 100 m.a.s.l; 120-140 m.a.s.l, 190-210 m.a.s.l) (Figure 1). Recently-emerged adults of *Z. fenestrata* Clade 1 were collected from under stones in rapids or in the moss or vegetation next to the stream and immediately stored in absolute ethanol. Large nymphs were also collected from under stones in rapids and returned to the laboratory in a cooler, where they were reared in Styrofoam cups at 11°C in water from their natal stream with small amounts of stream vegetation. Upon emerging as adults (within 30 days of sampling), individuals were immediately transferred to ethanol and stored at 4°C. While the exact location was not identified for each sample, the approximate altitude was recorded within 20 m altitude. Samples from within a locality were obtained from numerous different rocks across each sampling location.

### MORPHOLOGICAL CLASSIFICATION

All 127 individuals collected were photographed using a stereo microscope, and forewing length and body length were measured from a stage micrometer scale in ImageJ ^73^. Forewings and hindwings are equally sized for each individual, therefore measuring both was not necessary. We visually sorted specimens into either a fully winged (macropterous) or vestigial-winged (micropterous) groups. To examine the variation in wing length and body length we then visualized these data, and created a generalized linear model (GLM) for wing length based on body length, sex, sampling altitude and our previous wing length classification in R. These analyses tested for a clear pattern of wing dimorphism in this population, and to ensure the morphology classification was not biased by any additional influencing factors (e.g. size, altitude or sex).

### DNA EXTRACTION AND SEQUENCING

DNA extractions and GBS library prep were carried for 96 individuals (34 fully winged, 62 vestigial-winged) using the same methodology as Dussex, et al. ^63^. DNA extractions were carried out using DNeasy kits (Qiagen, Valencia, CA, USA) according to the manufacturer’s protocol using dissected head and femur tissue. Genotyping by sequencing library preparation followed the protocols of Elshire et al. (2011) with modifications as follows. DNA extractions were first dried using a vacuum centrifuge at 45°C, then resuspended in 15 μL dH2O. To each sample, a uniquely barcoded PstI adapter was added (2.25 ng per sample; Morris et al. 2011). DNA digestion was performed using 4UPstI-HF (NewEngland Biolabs, Ipswich, MA; Morris et al. 2011) in 1X CutSmart BufferTM130 with incubation at 37°C for 2 h. Adapters were ligated with T4 DNA ligase in 1X ligation buffer (New England Biolabs), followed by incubation at 16°C for 90 min and 80°C for 30 min. Purification was performed using a Qiagen MinElute PCR purification kit, with elution in 25 mL 1X TE. PCRs were carried out in 50 mL volumes containing 10 mL purified DNA, 1X MyTaqTM HS Master Mix (Bioline), and 1 mM each of PCR primers

5_AATGATACGGCGACCACCGAGATCTACACTCTTTCCCTACACGACGCTC TTCCGATC*T and 5_ CAAGCAGAAGACGGCATACGAGATCGGTCTCGGCATTCCTGCTGAACCGC TCTTCCGATC*T (where * indicates phosphorothioation) as per Dussex et al. (2016). PCRs were run in a Mastercycler ep Gradient S (Eppendorf, Hamburg, Germany) under the following conditions: 72°C for 5 min, 95°C for 60 s, and 24 cycles of 95°C for 30 s, 65°C for 30 s, and 72°C for 30s, with a final extension step at 72°C for 5 min. Sample concentrations were assessed using a NanoDrop spectrophotometer (Thermo Scientific) and all samples were pooled (20 ng DNA per sample). Size fractionation of the pooled library was achieved via electrophoresis on a 1.5% agarose gel, with a 300 bp size range from 200 to 500 bp selected for sequencing. A total of 96 samples were sequenced on one lane of an Illumina HiSeq 2500.

## ANALYSES

### Bioinformatic processing

All reads were trimmed, filtered and analyzed using the STACKS pipeline ^74^ in order to create catalogues of comparable SNP loci. We optimized the pipeline according to the recommendations of Paris, et al. ^75^. Initially, the PROCESS_RADTAGS module was used to separate reads by their barcode, remove low-quality reads (any read with an average Phred score < 10 in any sliding window of 11bp), trim all reads to 70 base pairs in length, and remove any reads that did not contain the enzyme recognition sequence. Next, the USTACKS module was used for the *de novo* assembly of raw reads into RAD tags. The minimum number of reads to create a stack was set at 3 (-m parameter in USTACKS), and the maximum number of pairwise differences between stacks was 2 (-M parameter in USTACKS). A catalogue of RAD tags was then generated using the 25 highest coverage individuals from each ecotype in CSTACKS. The distance allowed between catalogue loci (-n in CSTACKS) was increased to 2, after different trials were run to ensure loci were not inaccurately called as separate stacks. The execution of these components was accomplished using the STACKS denovo_map.pl script; in running this script, the optional -t flag was used to remove highly repetitive RAD tags during the USTACKS component of the pipeline. Following assembly and genotyping, the data were further filtered to maximize data quality. Using the POPULATIONS module, we retained only those loci that were genotyped in ≥50% of individuals and had a minor allele frequency ≥0.05 and a minimum stack depth of 10 (-m in POPULATIONS) for each individual. Genotypic data were exported from STACKS in GENEPOP format ^76^ and converted for subsequent analyses using PGD SPIDER v. 2 ^77^.

### Population Structure

We investigated the number of populations (or clusters) represented in our data using FASTSTRUCTURE ^78^ and the putatively neutral SNP dataset, default parameters, a logistic prior, and *K* from 1 to 6. The appropriate number of model components that explained structure in the dataset was determined using the *chooseK.py* function ^78^. Results for the identified optimal values of *K* were visualized using DISTRUCT ^79^. We also estimated the number of clusters using the *find.clusters* command in ADEGENET, with optimization based on the Bayesian Information Criterion (BIC). Finally, we created a Euclidian distance matrix between individuals in the R package ADEGENET ^80^, which we then displayed using a neighbor-joining tree produced in the R package APE ^81^.

### Outlier loci detection and annotation

Due to the limitations of differentiation-based methods and the potentially high false positive rates when looking for outlier loci under divergent selection ^82,83^, we utilized two distinct approaches: 1) an *FST* based outlier approach between *a priori* morphotype-pairs implemented in BAYESCAN ^84^; and 2) a hierarchical Bayesian modeling approach implemented in PCADAPT ^85^.

BAYESCAN analyses can give spurious results when there is significant over-representation of one of the groups being compared ^86^. Due to the sample size of vestigial-winged specimens being approximately twice the number of fully winged specimens, we performed two independent BAYESCAN runs, both including all fully winged individuals, but each with a different half of the vestigial-winged group. These two comparisons therefore each had a balanced design, and can be used to evaluate the generality of outlier loci detected across partially independent comparisons (given that one comparison group remains the same while the other changes). For each analysis, BAYESCAN was run using 10,000 output iterations, a thinning interval of 10, 20 pilot runs of length 10,000, and a burn-in period of 10,000, with prior odds of the neutral model of 10. We recorded all loci with a q-value of 0.2 or less, which equates to a false discovery rate of 20%. Q-values are far more stringent than p-values in classical statistics as they are adjusted for the false discovery rate given multiple comparisons, rather than the individual false positive rates in each comparison ^87^. To better understand the rates of false positive identification for outlier loci in this dataset, we also undertook 20 runs of BAYESCAN using identical parameters but comparing randomized groups of individuals (each also consisting of 34 individuals).

We also conducted outlier detection as implemented in PCADAPT ^85^. The number of Principal Components retained (*K*) for each analysis was determined by the graphical approach based on the scree-plot ^88^, as recommended by Luu, et al. ^85^.

## Results

### Morphology

Of 127 adults measured in this *Z. fenestrata* Clade 1 population, we found clear wing dimorphism for both males and females, with an approximately even number of each sex sampled (Figure 2). Fully winged individuals had an average forewing length: body length ratio of 1.06 ± 0.15, while the vestigial-winged individuals had an average forewing length: body length ratio of 0.26 ± 0.28, and there was no overlap in the distribution of wing lengths between groups. This difference in wing length was highly significant (t = −57.479, p < 2e-16). Sampling altitude (over this small altitudinal range) had no significant effect on the proportion of each morphotype, nor did it affect body length or wing length. Sex was significantly correlated with forewing length (t =-3.331, p = 0.00114), with females consistently having both longer forewings and bodies than males for both the fully winged and vestigial-winged forms, and there was also a significant positive correlation between body length and wing length within each sex (t = 2.811, p = 0.00575).

**Figure 2.**
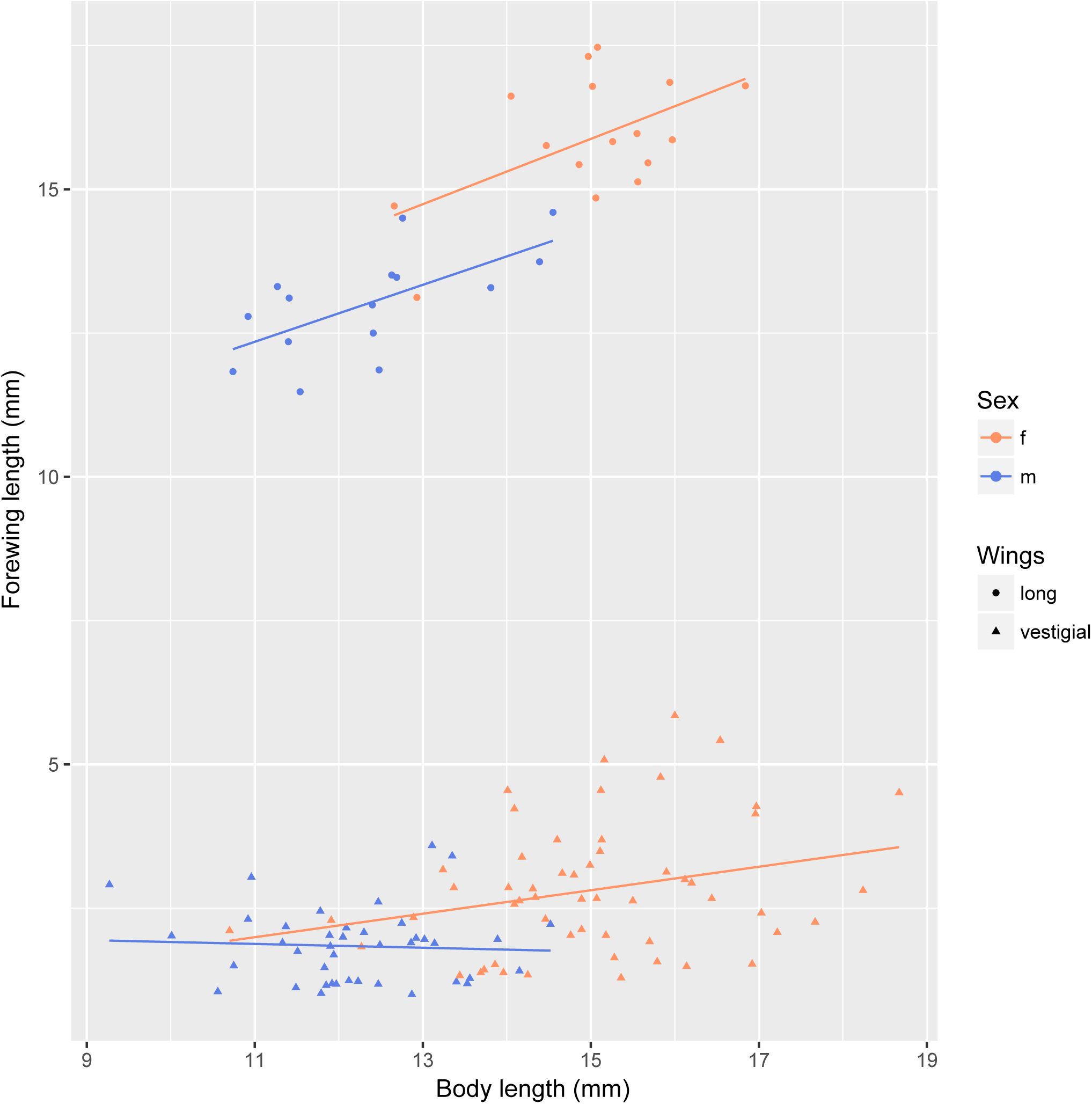
Variation in the relative wing length and body length of *Z. fenestrata* Clade 1 from Black Jacks Creek.

### GBS genotypic data and alignment

Following GBS, processing and filtering, we collected genotypic data at 6,843 SNPs across 96 of the measured 127 *Z. fenestrata* Clade 1 individuals – leaving out randomly selected vestigial-winged individuals as this dataset was far larger than the fully winged dataset. The sequences of these tags containing these SNPS are provided in Supplementary Table 1.

We found no detectable population structure across the samples using any of the analyses. FASTSTRUCTURE indicated an optimal number of clusters as 1, and when the higher number of clusters were investigated no clear pattern of differentiation emerged (Supplementary table 1). Similarly, using the *find.clusters* function in ADEGENET, the optimal number of clusters was 1, and no trend in differential clustering was visible for higher values of K. Finally, no genetic structure was evident in the neighbor-joining tree (Figure 3) or principal component analyses (Figure 4).

**Figure 3.**
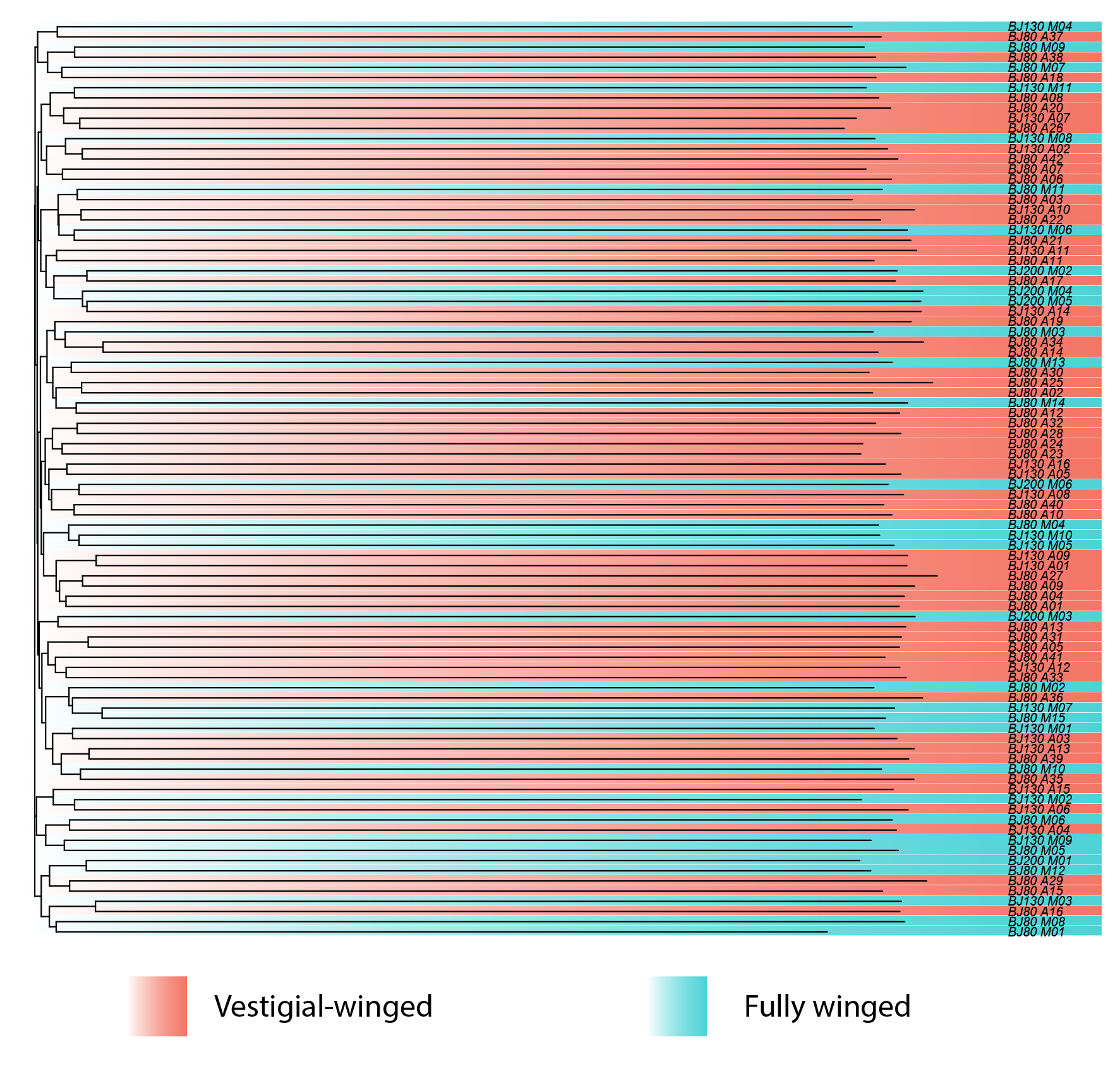
Neighbor-joining tree of *Z. fenestrata* Clade 1 samples showing the lack of phylogenetic differentiation between wing morphotypes

**Figure 4.**
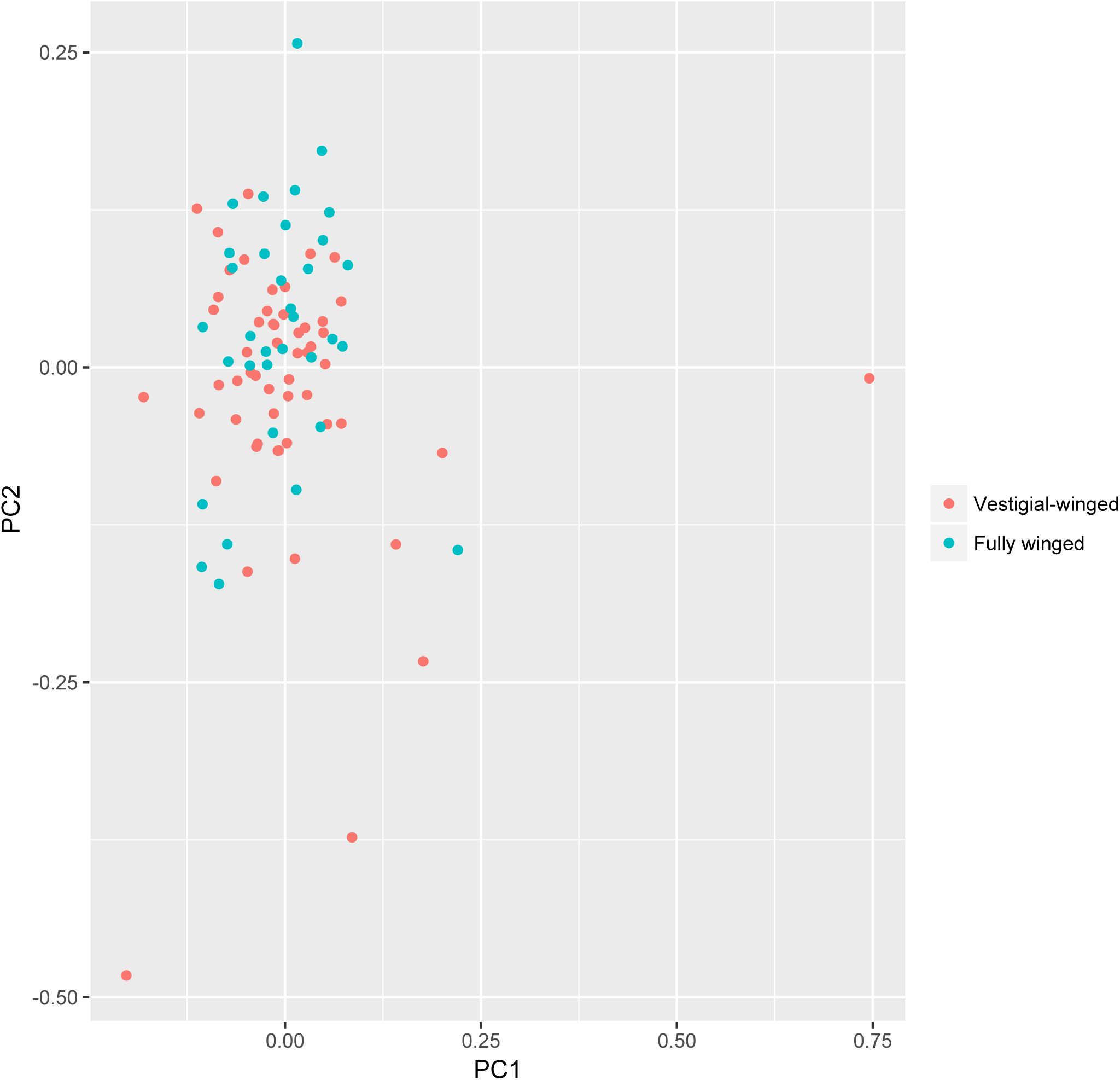
Principal component analysis of *Z. fenestrata* Clade 1 genetic differentiation in Black Jacks Creek.

**Figure 5.**
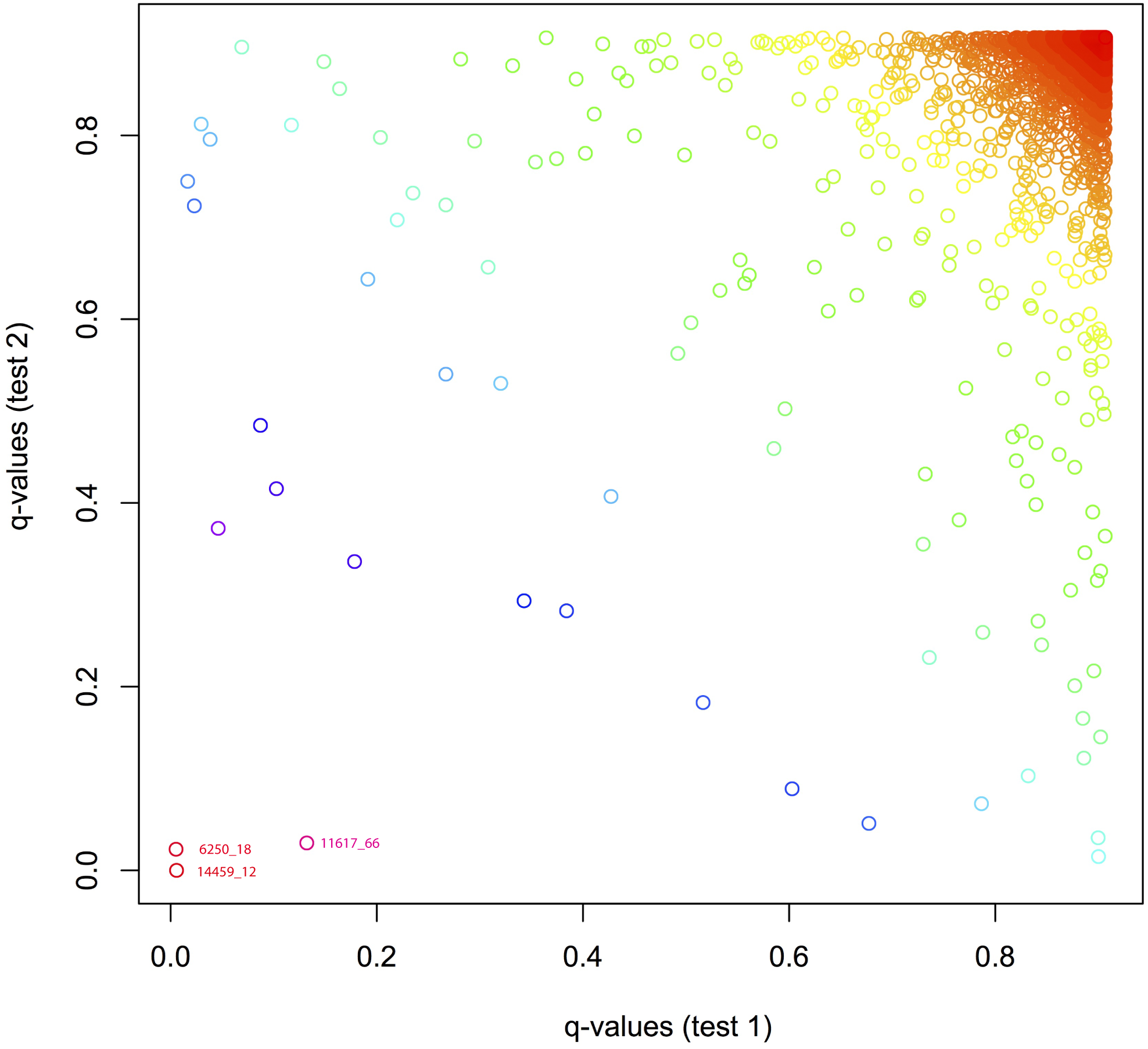
Scatterplot comparing the q-values obtained from the two independent BAYESCAN comparisons of fully winged and vestigial-winged morphotypes of *Z. fenestrata* Clade 1 sampled in Black Jacks Creek.

Given these results we conclude that there is no neutral population structure between fully winged and vestigial-winged individuals when sampled from the same location, and no differentiation among sampling localities. Given this apparent panmixia, genetic differences associated with morphotype differentiation, if present, must therefore be limited to small regions of the genome, likely indicating loci under divergent selection.

### Outlier loci detection and comparison

Given that no principal components correlated to morphotype differentiation, PCADAPT was unable to detect outliers associated with morphotypes, instead only identifying loci associated with the differentiation of a handful of slightly divergent individuals (Figure 4).

Because we had 34 fully winged individuals compared with 62 vestigial-winged individuals, we conducted two separate BAYESCAN analyses, dividing the vestigial-winged population sample in two. This was done because having highly uneven sample sizes in the two groups can disproportionately skew results ^86^. This approach also gave us the opportunity to compare the results of these two analyses, identifying loci that were found to be significant in these largely independent comparisons.

The two BAYESCAN runs detected 17 and 14 outlier loci with a q-value of <0.2 (Supplementary Table 2). Of these, three loci were identified in both comparisons, with one locus (14459_12) identified as the most significantly differentiated SNP in both comparisons, with q-values of (0.00570 and <0.00000). In independent comparisons with random differences between groups with loci differentiation distributions to those observed, one would expect 0.03 loci to be detected as outliers in both comparisons, and the probability that the most differentiated locus would be identical would be <0.0001.

In the randomized BAYESCAN runs, an average of 10.6 outlier loci were detected at a q-value of 0.2, with a maximum of 13 outlier loci detected. This number of outliers recorded is slightly lower than the real winged vs. wingless comparisons, however not greatly, indicating that at this relatively relaxed reporting value for q-values many of the recorded outliers are likely to be false positives. However, the minimum q-value recorded across these random comparisons was 0.026. In both of our real comparisons between winged and wingless groups, three outliers were more significant than this, including the outliers identified in multiple comparisons which were considerably lower. This provides strong evidence that these very high confidence outliers are truly associated with the different in phenotype and not statistical false positives.

The observed differentiation between fully winged and vestigial-winged individuals at these outlier loci strongly suggests that there are regions of the genome highly differentiated between these two morphotypes. Due to the paucity of genomic data published for Plecoptera, we were unable to map these outlier loci via BLAST-n to genomic regions to identify the genes present in the surrounding regions.

## Discussion

In this study, we tested for a genetic basis for wing reduction in the New Zealand stonefly *Z. fenestrata* Clade 1. While we found no neutral population structure among the two sympatric morphotypes we detected outlier loci between fully winged and vestigial-winged *Z. fenestrata* Clade 1 individuals, with several of the most highly differentiated outlier loci common to distinct sample comparisons. These results match the predictions of a ‘divergence with gene flow’ scenario, where small regions of the genome (genomic islands of divergence) are highly differentiated, contrasting with lower differentiation across the rest of the genome ^89-91^. These results strongly support the hypothesis that wing reduction in *Z. fenestrata* Clade 1 is at least partially genetically determined, and not solely an environmentally determined polyphenism.

Given a probable genetic basis for wing morphotype, and evidence for divergent selection for different morphotypes at different altitudes as indicated by the broader altitudinal distribution of the two morphotypes ^32,63^, this system is potentially an example of early ecological divergence with gene flow, similar to recent examples of ecological speciation e.g. ^92,93^. While reproductive barriers do not apparently exist between these two sympatric morphotypes in Clade 1, the broad system we describe demonstrates the effects of divergent selection at different altitudes, with ongoing gene flow where the two forms meet.

When populations occupy different habitats, divergent natural selection can cause differentiation in ecologically important characters (for review, see Schluter ^94^), and conversely, gene flow between divergent populations acts as a homogenizing force, eroding population differentiation ^95^. In the majority of *Z. fenestrata* Clade 1 populations, vestigial-winged populations occupy higher altitudes and are largely allopatric to the lower altitude fully winged populations. It appears that gene flow over any distance is extremely low for *Z. fenestrata,* as evidenced by the fine-scale genetic structure between nearby streams ^63^. This poor flighted dispersal ability may contribute towards maintaining the divergence between morphotype populations, despite the observed homogenization across the majority of the genome in geographic regions of population overlap. Indeed, the micropterous phenotype is likely to decrease gene flow due to the lack of any flighted long-distance dispersal. In most systems where ecological divergence is detected there is considerable reproductive isolation between morphotypes; the low dispersal abilities of *Z. fenestrata* may be the mechanism that helps maintain this isolation in most streams.

One question that remains to be addressed is why the Black Jacks Creek *Z. fenestrata* Clade 1 population exhibits the high degree of overlap between morphotypes, particularly relating to high proportion of vestigial-winged individuals present at low altitudes. Previous studies have indicated a sharp transition from fully winged to vestigial-winged or apterous at around 500 m.a.s.l. ^32^. We offer two hypotheses as to why sympatry occurs at this altitude at Black Jacks Creek, though these must be regarded as speculation until further testing is done. Firstly, a disturbance such as a large storm may have flushed out a large proportion of the fully winged individuals into the nearby Clutha River, replacing them with vestigial-winged individuals from higher altitudes. Alternatively, the selection pressure for wing reduction occurs at a lower altitude in this stream – or relates to very fine-scale microhabitat surrounding Black Jacks Creek, which is a patchy mosaic of scrub and grassland modified by recent farming activities.

Our results reinforce the need for taxonomic revision for this species group, as there is no genetic evidence for the separation of vestigial-winged morphotypes into the separate taxon *Z. pennulata.* Along with there being no neutral genetic differentiation between co-occurring morphotypes of this species, we found no temporal or spatial segregation of the two morphotypes, given that recently-emerged fully winged and vestigial-winged individuals were collected simultaneously. These results are consistent with the completely overlapping temporal patterns of emergence documented by McLellan ^62^.

While we infer that there is evidence for a genetic component to the differentiation of wing morphotypes, there may also be an environmental component to this differentiation. In other species of insects, the penetrance of genetic factors regulating wing development can be mediated by environmental factors, and therefore the expression of phenotype can be highly complex ^71,72^. The differing patterns of wing loss in the different clades of *Z. fenestrata* Clade 1 may indicate the interactive roles played between the environment and genetics. It remains possible that some level of environmentally determined gene expression is partially responsible for the observed wing morphotypes found across the *Z. fenestrata* species group.

While we analysed SNP data, we do not infer that SNPs underlie the phenotypic differences observed, nor that the outlier SNPs identified in our study have any causal relationship to the observed developmental differences between morphotypes. Rather, these SNPs are likely to be in linkage with changes in nearby regions of the genome that influence morphotype ^96^. As regions linked to the genetic changes underlying phenotypic differences can be very large (e.g. ^97,98^ we would require a well annotated and near complete genomic sequence before we could speculate as to the specific changes responsible for wing polymorphism.

Untangling the precise mechanisms behind wing reduction in the *Z. fenestrata* species group, including testing for an environmentally induced component to these alternative developmental pathways will require further experimentation. While the *Z. fenestrata* species group is a fascinating system to study the mechanisms wing reduction in insects, the group does have some life-history and population characteristics that create challenges for understanding the mechanism(s) behind wing loss difficult. *Z. fenestrata* can have a long generation time (perhaps involving years as a wingless nymph), making breeding experiments and QTL studies challenging.

Furthermore, their habitat is fast flowing rapids in highly oxygenated streams with cold water, making them difficult to raise in laboratory settings for a full life cycle, and hindering reciprocal translocation experiments in the wild. Combining long-term common garden experiments and analyses of gene expression should provide more information to the regulatory mechanisms and pathways for wing development in this species.

Currently the genomic resources for *Z. fenestrata* (and all Plecoptera) are too incomplete to determine if the outlier loci identified are adjacent to each other, or more generally, if they are in islands of divergence. Without these genomic resources, it is also impossible to speculate as to the potential underlying genes that may be responsible for these two phenotypes. With further work creating a genome assembly for this species we will be able to look at the specific genomic regions linked to the outlier SNPs defined in this study.

## Conclusion

Wing dimorphism is a common trait across many species of stoneflies, but the mechanisms behind this have yet to be investigated. *Z. fenestrata* Clade 1 presents an ideal taxon to examine this, potentially revealing the generalized mechanisms behind wing reduction in this order. Our results for this spatially overlapping population of fully winged and vestigial-winged *Z. fenestrata* Clade 1 morphotypes supports the hypothesis that wing development has a genetic mechanism rather than being solely environmentally determined. While there was no neutral genetic structure between wing morphotypes, outlier loci were identified between these two groups. While it is possible that these outlier loci are not themselves linked with the specific causative changes associated with wing development, any genetic differences linked to wing morphotype differentiation in an otherwise sympatric population must indicate that there is some genetic differentiation between morphotypes. Further examination of these outlier loci may reveal the underlying genes linked to wing reduction in this species.

## Acknowledgements

We wish to thank Tania King for her assistance in the lab, and Maxim Nekrasov at the John Curtin School of Medicine for his assistance in library sequencing. Thanks to Graham McCulloch and Brian Patrick for their advice on the system. This work was funded by Marsden fund grant (UOO1412).

## Author contributions statement

AV planned and wrote the manuscript and performed the analyses, BF and JW conducted the fieldwork, JW and PD envisioned and planned the project, all authors edited and redrafted the manuscript.

## Data Accessibility

All processed data from Stacks will be included on Dryad entry # XXXXXXX

## Ethical Statement

All experiments were performed in accordance with University of Otago ethics committee regulations and guidelines.

## References

1 Harrison, R. G. DISPERSAL POLYMORPHISMS IN INSECTS. Annual Review of Ecology and Systematics 11, 95–118, doi:10.1146/annurev.es.11.110180.000523 (1980).

2 Roff, D. A. THE EVOLUTION OF WING DIMORPHISM IN INSECTS. Evolution 40, 1009–1020, doi:10.2307/2408759 (1986).

3 Masaki, S. & Shimizu, T. VARIABILITY IN WING FORM OF CRICKETS. Researches on Population Ecology 37, 119–128, doi:10.1007/bf02515769 (1995).

4 Zera, A. J. & Denno, R. F. Physiology and ecology of dispersal polymorphism in insects. Annual Review of Entomology 42, 207–230, doi:10.1146/annurev.ento.42.1.207 (1997).

5 Zera, A. J. The endocrine regulation of wing polymorphism in insects: State of the art, recent surprises, and future directions. Integrative and Comparative Biology 43, 607–616, doi:10.1093/icb/43.5.607 (2003).

6 Roff, D. A. THE EVOLUTION OF FLIGHTLESSNESS - IS HISTORY IMPORTANT. Evolutionary Ecology 8, 639–657, doi:10.1007/bf01237847 (1994).

7 Roff, D. A. THE EVOLUTION OF FLIGHTLESSNESS IN INSECTS. Ecological Monographs 60, 389–421, doi:10.2307/1943013 (1990).

8 Roff, D. A. & Fairbairn, D. J. WING DIMORPHISMS AND THE EVOLUTION OF MIGRATORY POLYMORPHISMS AMONG THE INSECTA. American Zoologist 31, 243–251 (1991).

9 Roff, D. A. HABITAT PERSISTENCE AND THE EVOLUTION OF WING DIMORPHISM IN INSECTS. American Naturalist 144, 772–798, doi:10.1086/285706 (1994).

10 Wagner, D. L. & Liebherr, J. K. FLIGHTLESSNESS IN INSECTS. Trends Ecol. Evol. 7, 216–220, doi:10.1016/0169-5347(92)90047-f (1992).

11 Denno, R. F., Hawthorne, D. J., Thorne, B. L. & Gratton, C. Reduced flight capability in British Virgin Island populations of a wing-dimorphic insect: the role of habitat isolation, persistence, and structure. Ecological Entomology 26, 25–36, doi:10.1046/j.1365-2311.2001.00293.x (2001).

12 Den Boer, P. J. On the significance of dispersal power for populations of carabid-beetles (Coleoptera, Carabidae). Oecologia 4, 1–28, doi:10.1007/bf00390612 (1970).

13 Brinck, P. Studies on Swedish stoneflies. Opuscula Entomologica 11 11, 1–250 (1949).

14 Jackson, D. J. The inheritance of long and short wings in the weevil, Sitonia hispidula, with a discussion of wing reduction among beetles. Transactions of the Royal Society of Edinburgh 55, 655–735 (1928).

15 Hynes, H. B. N. The taxonomy and ecology of the nymphs of British Plecoptera with notes on the adults and eggs. Trans Roy Ent Soc London 91, 459–557 (1941).

16 Roff, D. A. THE COST OF BEING ABLE TO FLY – A STUDY OF WING POLYMORPHISM IN 2 SPECIES OF CRICKETS. Oecologia 63, 30–37, doi:10.1007/bf00379781 (1984).

17 Zera, A. J. DIFFERENCES IN SURVIVORSHIP, DEVELOPMENT RATE AND FERTILITY BETWEEN THE LONGWINGED AND WINGLESS MORPHS OF THE WATERSTRIDER, LIMNOPORUS-CANALICULATUS. Evolution 38, 1023–1032, doi:10.2307/2408436 (1984).

18 Roff, D. A. & Bradford, M. J. Quantitative genetics of the trade-off between fecundity and wing dimorphism in the cricket Allonemobius socius. Heredity 76, 178–185, doi:10.1038/hdy. 1996.25 (1996).

19 Roff, D. A., Tucker, J., Stirling, G. & Fairbairn, D. J. The evolution of threshold traits: effects of selection on fecundity and correlated response in wing dimorphism in the sand cricket. Journal of Evolutionary Biology 12, 535–546 (1999).

20 Ikeda, H., Kagaya, T., Kubota, K. & Abe, T. Evolutionary relationships among food habit, loss of flight, and reproductive traits: Life-history evolution in the Silphinae (Coleoptera : Silphidae). Evolution 62, 2065–2079, doi:10.1111/j.1558-5646.2008.00432.x (2008).

21 Langellotto, G. A., Denno, R. F. & Ott, J. R. A trade-off between flight capability and reproduction in males of a wing-dimorphic insect. Ecology 81, 865–875, doi:10.1890/0012-9658(2000)081[0865:atobfc]2.0.co;2 (2000).

22 Thomas, M. A., Walsh, K. A., Wolf, M. R., McPheron, B. A. & Marden, J. H. Molecular phylogenetic analysis of evolutionary trends in stonefly wing structure and locomotor behavior. Proceedings of the National Academy of Sciences of the United States of America 97, 13178–13183, doi:10.1073/pnas.230296997 (2000).

23 Marden, J. H. & Kramer, M. G. SURFACE-SKIMMING STONEFLIES – A POSSIBLE INTERMEDIATE STAGE IN INSECT FLIGHT EVOLUTION. Science 266, 427–430, doi:10.1126/science.266.5184.427 (1994).

24 Thomas, A. L. R. & Norberg, R. A. Skimming the surface – The origin of flight in insects? Trends Ecol. Evol. 11, 187–188, doi:10.1016/0169-5347(96)30022-0 (1996).

25 Samways, M. J. Skimming and insect evolution. Trends Ecol. Evol. 11, 471471, doi:10.1016/0169-5347(96)81156-6 (1996).

26 Marden, J. H. & Thomas, M. A. Rowing locomotion by a stonefly that possesses the ancestral pterygote condition of co-occurring wings and abdominal gills. Biol. J. Linnean Soc. 79, 341–349, doi:10.1046/j.1095-8312.2003.00192.x (2003).

27 Costello, M. J. Preliminary Observations on Wing-Length Polymorphism in Stoneflies (Plecoptera: Insecta). The Irish Naturalists' Journal 22, 474–478 (1988).

28 Brundin, L. INSECTS AND PROBLEM OF AUSTRAL DISJUNCTIVE DISTRIBUTION. Annual Review of Entomology 12, 149-&, doi:10.1146/annurev.en.12.010167.001053 (1967).

29 Zwick, P. Phylogenetic system and zoogeography of the plecoptera. Annual Review of Entomology 45, 709–746, doi:10.1146/annurev.ento.45.1.709 (2000).

30 Schultheis, A. S., Weigt, L. A. & Hendricks, A. C. Gene flow, dispersal, and nested clade analysis among populations of the stonefly Peltoperla tarteri in the southern Appalachians. Mol. Ecol. 11, 317–327, doi:10.1046/j.1365-294X.2002.01445.x (2002).

31 Fochetti, R. & de Figueroa, J. M. T. Global diversity of stoneflies (Plecoptera : Insecta) in freshwater. Hydrobiologia 595, 365–377, doi:10.1007/s10750-007-9031-3 (2008).

32 McCulloch, G. A., Wallis, G. P. & Waters, J. M. Do insects lose flight before they lose their wings? Population genetic structure in subalpine stoneflies. Mol. Ecol. 18, 4073–4087, doi:10.1111/j.1365-294X.2009.04337.x (2009).

33 McCulloch, G. A., Wallis, G. P. & Waters, J. M. A time-calibrated phylogeny of southern hemisphere stoneflies: Testing for Gondwanan origins. Mol. Phylogenet. Evol. 96, 150–160, doi:10.1016/j.ympev.2015.10.028 (2016).

34 Lillehammer, A. NORWEGIAN STONE-FLIES PART 5 VARIATIONS IN MORPHOLOGICAL CHARACTERS COMPARED TO DIFFERENCES IN ECOLOGICAL FACTORS. Norwegian Journal of Entomology 23, 161–172 (1976).

35 Malmqvist, B. How does wing length relate to distribution patterns of stoneflies (Plecoptera) and mayflies (Ephemeroptera)? Biol. Conserv. 93, 271–276, doi:10.1016/s0006-3207(99)00139-1 (2000).

36 Loskutova, O. A. & Zhiltzova, L. A. Wing and body size polymorphism in populations of the stonefly Arcynopteryx dichroa McL. (Plecoptera: Perlodidae) in the Ural Mountains, Russia. Polar Research 35, doi: 10.3402/polar.v35.26596 (2016).

37 Saltveit, S. J. & Brittain, J. E. Short-wingedness in the stonefly Diura nanseni (Kempny) (Plecoptera: Perlodidae). Entomologica Scandinavica 17, 153–156 (1986).

38 Westermann, F. WING POLYMORPHISM IN CAPNIA-BIFRONS (PLECOPTERA, CAPNIIDAE). Aquatic Insects 15, 135–140, doi:10.1080/01650429309361510 (1993).

39 Donald, D. B. & Patriquin, D. E. THE WING LENGTH OF LENTIC CAPNIIDAE (PLECOPTERA) AND ITS RELATIONSHIP TO ELEVATION AND WISCONSIN GLACIATION. Canadian Entomologist 115, 921–926 (1983).

40 Davey, J. W. et al. Genome-wide genetic marker discovery and genotyping using next-generation sequencing. Nature Reviews Genetics 12, 499–510, doi:10.1038/nrg3012 (2011).

41 Narum, S. R., Buerkle, C. A., Davey, J. W., Miller, M. R. & Hohenlohe, P. A. Genotyping-by-sequencing in ecological and conservation genomics. Mol. Ecol. 22, 2841–2847, doi:10.1111/mec.12350 (2013).

42 Ellegren, H. Genome sequencing and population genomics in non-model organisms. Trends Ecol. Evol. 29, 51–63, doi:10.1016/j.tree.2013.09.008 (2014).

43 Andrews, K. R., Good, J. M., Miller, M. R., Luikart, G. & Hohenlohe, P. A. Harnessing the power of RADseq for ecological and evolutionary genomics. Nature Reviews Genetics 17, 81–92, doi:10.1038/nrg.2015.28 (2016).

44 Stinchcombe, J. R. & Hoekstra, H. E. Combining population genomics and quantitative genetics: finding the genes underlying ecologically important traits. Heredity 100, 158–170, doi:10.1038/sj.hdy.6800937 (2008).

45 Stapley, J. et al. Adaptation genomics: the next generation. Trends Ecol. Evol. 25, 705–712, doi: 10.1016/j.tree.2010.09.002 (2010).

46 Pardo-Diaz, C., Salazar, C. & Jiggins, C. D. Towards the identification of the loci of adaptive evolution. Methods in Ecology and Evolution 6, 445–464, doi:10.1111/2041-210x. 12324 (2015).

47 Long, A. D. & Langley, C. H. The power of association studies to detect the contribution of candidate genetic loci to variation in complex traits. Genome Research 9, 720–731 (1999).

48 Shimizu, K. K. & Purugganan, M. D. Evolutionary and ecological genomics of arabidopsis. Plant Physiology 138, 578–584, doi:10.1104/pp.105.061655 (2005).

49 Stranger, B. E., Stahl, E. A. & Raj, T. Progress and Promise of Genome-Wide Association Studies for Human Complex Trait Genetics. Genetics 187, 367–383, doi:10.1534/genetics.110.120907 (2011).

50 Ogawa, K. & Miura, T. Two developmental switch points for the wing polymorphisms in the pea aphid Acyrthosiphon pisum. Evodevo 4, doi:10.1186/2041-9139-4-30 (2013).

51 Zera, A. J. Endocrine analysis in evolutionary-developmental studies of insect polymorphism: hormone manipulation versus direct measurement of hormonal regulators. Evolution & Development 9, 499–513 (2007).

52 Xu, H. J. et al. Two insulin receptors determine alternative wing morphs in planthoppers. Nature 519, 464-+, doi:10.1038/nature14286 (2015).

53 Lin, X. D., Yao, Y., Wang, B., Emlen, D. J. & Lavine, L. C. Ecological Trade-offs between Migration and Reproduction Are Mediated by the Nutrition-Sensitive Insulin-Signaling Pathway. International Journal of Biological Sciences 12, 607–616, doi:10.7150/ijbs.14802 (2016).

54 Abouheif, E. & Wray, G. A. Evolution of the gene network underlying wing polyphenism in ants. Science 297, 249–252, doi:10.1126/science.1071468 (2002).

55 Van Belleghem, S. M., Roelofs, D., Van Houdt, J. & Hendrickx, F. De novo Transcriptome Assembly and SNP Discovery in the Wing Polymorphic Salt Marsh Beetle Pogonus chalceus (Coleoptera, Carabidae). Plos One 7, doi: 10.1371/journal.pone.0042605 (2012).

56 Brisson, J. A. Aphid wing dimorphisms: linking environmental and genetic control of trait variation. Philosophical Transactions of the Royal Society B-Biological Sciences 365, 605–616, doi:10.1098/rstb.2009.0255 (2010).

57 Brisson, J. A., Ishikawa, A. & Miura, T. Wing development genes of the pea aphid and differential gene expression between winged and unwinged morphs. Insect Molecular Biology 19, 63–73, doi:10.1111/j.1365-2583.2009.00935.x (2010).

58 Nijhout, H. F. Control mechanisms of polyphenic development in insects – In polyphenic development, environmental factors alter same aspects of development in an orderly and predictable way. Bioscience 49, 181–192, doi:10.2307/1313508 (1999).

59 Zhou, X. S., Chen, J. L., Meizhang, Liang, S. K. & Wang, F. H. Differential DNA Methylation Between Two Wing Phenotypes Adults of Sogatella furcifera. Genesis 51, 819–826, doi:10.1002/dvg.22722 (2013).

60 Liang, S. K. et al. CpG methylated ribosomal RNA genes in relation to wing polymorphism in the rice pest Sogatella furcifera. Journal of Asia-Pacific Entomology 18, 471–475, doi:10.1016/j.aspen.2015.06.002 (2015).

61 McLellan, I. D. ALPINE AND SOUTHERN PLECOPTERA FROM NEW-ZEALAND, AND A NEW CLASSIFICATION OF GRIPOPTERYGIDAE. N. Z. J. Zool. 4, 119–147 (1977).

62 McLellan, I. D. A revision of Zelandoperla Tillyard (Plecoptera : Gripopterygidae : Zelandoperlinae). N. Z. J. Zool. 26, 199–219 (1999).

63 Dussex, N., Chuah, A. & Waters, J. M. Genome-wide SNPs reveal fine-scale differentiation among wingless alpine stonefly populations and introgression between winged and wingless forms. Evolution 70, 38–47, doi:10.1111/evo.12826 (2016).

64 Lindroth, C. H. Inheritance and wing dimorphism in Pterostichus anthracinus. Hereditas 32, 37–40 (1945).

65 Aukema, B. Wing-length determination in two wing-dimorphic Calathus species (Coleoptera: Carabidae). Hereditas 113, 189–202 (1990).

66 Harrison, R. G. Flight polymorphism in the field cricket Gryllus pennsylvanicus. Oecologia 40, 125–132 (1979).

67 Rose, D. J. W. Dispersal and quality in populations of Cicadulina species (Cicadellidae). J. Anim. Ecol. 41, 589–609 (1972).

68 Zera, A. J. & Tiebel, K. Differences in juvenile hormone esterase activity between presumptive macropterous and brachypterous Gryllus rubens: implications for the hormonal control of wing polymorphism. Journal of Insect Physiology 35, 7–17 (1989).

69 Kimura, T. & Masaki, S. Brachypterism and seasonal adaptation in Orgyia thyellina Butler (Lepidoptera, Lymantriidae). Kontyu 45, 97–106 (1977).

70 Vellichirammal, N. N., Madayiputhiya, N. & Brisson, J. A. The genomewide transcriptional response underlying the pea aphid wing polyphenism. Mol. Ecol. 25, 4146–4160, doi:10.1111/mec.13749 (2016).

71 Honek, A. FACTORS AND CONSEQUENCES OF A NONFUNCTIONAL ALARY POLYMORPHISM IN PYRRHOCORIS-APTERUS (HETEROPTERA, PYRRHOCORIDAE). Researches on Population Ecology 37, 111–118, doi:10.1007/bf02515768 (1995).

72 Vellichirammala, N. N., Guptab, P., Hallc, T. A. & Brisson, J. A. Ecdysone signaling underlies the pea aphid transgenerational wing polyphenism. Proceedings of the National Academy of Sciences of the United States of America 114, 1419–1423 (2017).

73 Schindelin, J., Arganda-Carreras, I. & Frise, E. Fiji: an open-source platform for biological-image analysis. Nature Methods 9, 676–682 (2012).

74 Catchen, J., Hohenlohe, P. A., Bassham, S., Amores, A. & Cresko, W. A. Stacks: an analysis tool set for population genomics. Mol. Ecol. 22, 3124–3140, doi:10.1111/mec.12354 (2013).

75 Paris, J. R., Stevens, J. R. & Catchen, J. Lost in parameter space: a road map for STACKS. Methods in Ecology and Evolution, doi: 10.1111/2041-210X.12775 (2017).

76 Raymond, M. & Rousset, F. Geneppop (Version 1.2) - Population-genetics software for exact tests and ecumenicism. J. Hered. 86 (1995).

77 Lischer, H. E. L. & Excoffier, L. PGDSpider: an automated data conversion tool for connecting population genetics and genomics programs. Bioinformatics 28, 298–299 (2012).

78 Raj, A., Stephens, M. & Pritchard, J. K. fastSTRUCTURE: Variational Inference of Population Structure in Large SNP Data Sets. Genetics 197, 573–U207, doi:10.1534/genetics. 114.164350 (2014).

79 Rosenberg, N. A. DISTRUCT: a program for the graphical display of population structure. Molecular Ecology Notes 4, 137–138, doi:10.1046/j.1471-8286.2003.00566.x (2004).

80 Jombart, T. adegenet: a R package for the multivariate analysis of genetic markers. Bioinformatics 24, 1403–1405, doi:10.1093/bioinformatics/btn129 (2008).

81 Paradis, E., Claude, J. & Strimmer, K. APE: Analyses of Phylogenetics and Evolution in R language. Bioinformatics 20, 289–290, doi:10.1093/bioinformatics/btg412 (2004).

82 De Mita, S. et al. Detecting selection along environmental gradients: analysis of eight methods and their effectiveness for outbreeding and selfing populations. Mol. Ecol. 22, 1383–1399 (2013).

83 Vilas, A., Perez-Figueroa, A. & Caballero, A. A simulation study on the performance of differentiation-based methods to detect selected loci using linked neutral markers. Journal of Evolutionary Biology 25, 1364–1376 (2012).

84 Foll, M. & Gaggiotti, O. A genome-scan method to identify selected loci appropriate for both dominant and codominant markers: a bayesian perspective. Genetics 180, 977–993, doi:10.1534/genetics.108.092221 (2008).

85 Luu, K., Bazin, E. & Blum, M. G. pcadapt: an R package to perform genome scans for selection based on principal component analysis. Molecular Ecology Resources 17, 67–77 (2017).

86 Helyar, S. J. et al. Application of SNPs for population genetics of nonmodel organisms: new opportunities and challenges. Mol. Ecol. Resour. 11, 123–136, doi:10.1111/j.1755-0998.2010.02943.x (2011).

87 Storey, J. D. & Tibshirani, R. Statistical significance for genomewide studies. Proceedings of the National Academy of Sciences of the United States of America 100, 9440–9445, doi:10.1073/pnas.1530509100 (2003).

88 Jackson, D. A. Stopping rules in principal components analysis : a comparison of heuristical and statistical approaches. Ecology, 2204–2214 (1993).

89 Lotterhos, K. E. & Whitlock, M. C. The relative power of genome scans to detect local adaptation depends on sampling design and statistical method. Mol. Ecol. 24, 1031–1046, doi:10.1111/mec.13100 (2015).

90 Nosil, P., Funk, D. J. & Ortiz-Barrientos, D. Divergent selection and heterogeneous genomic divergence. Mol. Ecol. 18, 375–402, doi:10.1111/j.1365-294X.2008.03946.x (2009).

91 Via, S. & West, J. The genetic mosaic suggests a new role for hitchhiking in ecological speciation. Mol. Ecol. 17, 4334–4345, doi: 10.1111/j.1365-294X.2008.03921.x (2008).

92 Schluter, D. & Conte, G. L. Genetics and ecological speciation. Proceedings of the National Academy of Sciences of the United States of America 106, 9955–9962, doi:10.1073/pnas.0901264106 (2009).

93 Nosil, P. Ecological Speciation. (Oxford University Press, 2012).

94 Schluter, D. The Ecology of Adaptive Radiation. (Oxford University Press, 2000).

95 Slatkin, M. Gene flow and the geographic structure of natural populations. Science 236, 787–792 (1987).

96 Visscher, P. M. et al. 10 Years of GWAS Discovery: Biology, Function, and Translation. American Journal of Human Genetics 101, 5–22 (2017).

97 Boitard, S., Boussaha, M., Capitan, A., Rocha, D. & Servin, B. Uncovering Adaptation from Sequence Data: Lessons from Genome Resequencing of Four Cattle Breeds. Genetics 203, 433-+, doi:10.1534/genetics.115.181594 (2016).

98 Schlamp, F. et al. Evaluating the performance of selection scans to detect selective sweeps in domestic dogs. Mol. Ecol. 25, 342–356, doi:10.1111/mec.13485 (2016).

